# Loss of microglia reduces NGF signaling and retinal ganglion cell survival

**DOI:** 10.64898/2026.03.26.714400

**Authors:** Lucia Buccarello, Giorgia Ribbeni, Laura Riccieri, Ophelia Livero, Antonino Cattaneo, Silvia Marinelli

## Abstract

Nerve growth factor (NGF) exerts neuroprotective effects in the retina, and accumulating evidence indicates that microglia represent a key cellular target of NGF/TrkA signaling. However, evidence showing that the NGF/TrkA signaling in microglia is required for downstream neuroprotective actions remains unresolved. Here, we directly addressed this question by pharmacologically depleting microglia and assessing the impact on NGF pathway activity and retinal integrity.

Adult C57BL/6J mice were treated with the CSF1R inhibitor PLX5622 for three weeks, resulting in a robust (∼77%) depletion of retinal microglia. Microglial ablation induced marked structural and cellular alterations, including significant loss of retinal ganglion cells (RGCs) and thinning of retinal layers, in the absence of any other lesion or insult. Residual microglia exhibited layer-specific phenotypic changes, with a phagocytic profile in the ganglion cell layer and a more ramified morphology in the outer plexiform layer.

Strikingly, microglial depletion led to a profound decrease of NGF signaling, with a strong reduction in total and phosphorylated TrkA, and decreased p75NTR levels, in retinal extracts. The amount of TrkA expression is strongly correlated with microglial levels, supporting a primary role of microglia in sustaining NGF signaling in the retina. Together, these findings demonstrate that microglia are required for NGF/TrkA signaling and identify these cells as essential mediators of NGF-dependent neuroprotection in the retina.

## Introduction

Microglia are the principal resident innate immune cells of the central nervous system (CNS), including the retina, where they form a crucial component of the glial network. Since their initial identification by Pío del Río-Hortega ^1^, microglia are now recognized for diverse roles beyond immune surveillance during CNS development and homeostasis ^2^. These include involvement in vascularization, neurogenesis, neuronal survival and synaptic pruning ^3^. In the adult retina, microglia exhibit a highly ramified morphology and dynamic motility, mirroring their counterparts in other CNS regions ^4^. Given their broad functional repertoire, retinal microglia represent a promising therapeutic target in both ocular and CNS neurodegenerative disorders. Their development and maturation in the retina are tightly regulated by a range of transcription factors and cytokine receptors ^2,5–7^. Additionally, colony-stimulating factor 1 receptor (CSF1R) plays a critical role as a survival receptor expressed on microglia throughout development ^8–10^. Retinal microglia originate from two distinct waves of infiltration during development ^9,11,12^, which overlap with the differentiation of retinal neurons into five principal types: photoreceptors, bipolar cells, amacrine cells, horizontal cells and retinal ganglion cells (RGCs). These neurons are organized into structured cellular and synaptic layers. The ganglion cell layer (GCL) contains the somata of RGCs, whose axons transmit visual information to the brain, while the outer plexiform layer (OPL) hosts synapses between photoreceptors and the dendrites of bipolar and horizontal cells ^13–16^. Microglia constitute approximately 0.2% of the total retinal cell population and coexist with astrocytes and Müller glia ^17–19^. In the retina, the distribution of microglia is layer-specific: nearly 80% of these cells are localized to the inner plexiform layer (IPL) and ganglion cell layer (GCL), and around 20% reside in the outer plexiform layer (OPL), while the outer nuclear layer (ONL) is largely devoid^12,14,20^. Their density and morphology vary with developmental stage ^12^, and their morphological state reflects functional status.

In adult mice, ramified microglia are characteristic of homeostasis and surveillance, whereas amoeboid microglia are associated with a reactive microglia phenotype ^21–23^. As described extensively in the CNS, this morphological shift also occurs in the retina in response to injury or stress, leading to the emergence of reactive, amoeboid microglia ^24,25^.

Persistent activation of microglia has been implicated in exacerbating tissue damage in preclinical models of retinal degeneration ^26,27^. Both genetic ^27,28^ and pharmacological ^29–31^ depletion strategies have shown that chronic microglial activation is an hallmark of many retinal pathologies. Importantly, pharmacological targeting of overactive microglia can mitigate disease progression. PLX5622, a highly selective and bioavailable CSF1R inhibitor capable of crossing the blood-brain barrier ^32^, has proven effective in transiently depleting microglia and has been used to investigate their role in retinal diseases such as glaucoma, ocular hypertension, optic nerve injury, and diabetic retinopathy ^27–31^. These studies have revealed both detrimental and neuroprotective roles of microglia, highlighting the complex, context-dependent nature of their functions. Furthermore, while microglia depletion do not impact plasticity of visual circuitry ^29^ targeting specific microglia genes, such as the purinergic receptor P2Y12, prevents it ^33^. Therefore, targeting microglia-specific signaling pathways or depleting microglia completely have different impact on the physiology of neuronal circuit and on the variety of neurodegenerative conditions. Crucially, the precise mechanisms by which microglial depletion affects neurotrophic signaling, such as that involving nerve growth factor (NGF), remain largely unexplored in the retina. In our previous work, we demonstrated that microglia serve as a critical cellular target for nerve growth factor (NGF)-induced TrkA signaling, contributing to RGC protection, identifying human recombinant NGF mutant (hNGFp) as a potent therapeutic candidate for retinal degeneration ^34^. Building on these findings, the present study investigates the effect of microglial removal on retinal health and NGF signalling using PLX5622 supplied in the diet. Specifically, we assessed the effects of microglial ablation on RGCs survival, glial cell state, and NGF signaling. Our data demonstrate that microglial depletion alters NGF receptor expression, RGC number as well as overall retina thickness.

Notably, we provide the first evidence of a direct interaction between microglia and TrkA NGF receptor in the retina, establishing microglia as essential mediators of NGF-dependent neuroprotective signaling.

## Methods

### Animal Model and Experimental Design

All procedures involving experimental animals adhered to the ARVO Statement for the Use of Animals in Ophthalmic and Vision Research and were approved by the competent Institutional Animal Care and Use Committee (authorization no. 102/2021-PR).

Adult male and female C57BL/6J mice (8–10 weeks old; Jackson Laboratories, Bar Harbor, ME, USA) were utilized for all experimental procedures (n = 30). Animals were housed under standard laboratory conditions with ad libitum access to food and water and maintained on a standard rodent chow diet for a 7-day acclimation period. Following acclimation, mice were randomly assigned to one of two dietary groups: a control group receiving standard chow or a treatment group receiving chow supplemented with 1200 mg/kg of PLX5622 (PLX; an inhibitor of the colony-stimulating factor 1 receptor) for 21 days to induce microglial depletion, as described by Elmore ^35^. At the conclusion of the treatment period, animals were euthanized by cervical dislocation. Both eyes and optic nerves were promptly harvested to evaluate the effects of PLX5622 on microglial and neuronal populations. Enucleated eyes were stored in phosphate-buffered saline (PBS) at 4 °C, and retinal dissections were performed immediately post-mortem. For ex-vivo analysis, whole eyes were allocated for either immunofluorescence analysis (n = 9 per group) or biochemical assays (n = 5 per group).

### Preparation of retinal lysates

Retinae and optic nerves were harvested and processed separately. Each sample was lysed in 93 μL of lysis buffer composed of 1% Triton X-100 (Sigma-Aldrich, Milan, Italy; Cat. No. 9002-93-1), a protease inhibitor cocktail (Complete, Roche Diagnostics, Basel, Switzerland), and phosphatase inhibitors (Sigma-Aldrich, St. Louis, MO, USA). The lysis buffer also contained the following components (in mM): 20 TRIS-acetate, 0.27 sucrose, 1 EDTA, 1 EGTA, 1 sodium orthovanadate, 50 sodium fluoride, 5 sodium pyrophosphate, 10 sodium β-glycerophosphate, and 1 dithiothreitol (DTT) (all from Sigma-Aldrich, Milan, Italy). Samples were incubated on ice for 30 minutes to facilitate protein extraction, followed by centrifugation at 10,000 rpm for 10 minutes at 4 °C. The supernatants were collected and stored at −20 °C until further use.

### Western Blot Analysis

Protein concentrations were determined using the Bradford assay (Bio-Rad Protein Assay; Cat. No. 500-0006, Munich, Germany). Equal amounts of protein (40 μg) were separated by 12% SDS-polyacrylamide gel electrophoresis (SDS-PAGE) and transferred onto PVDF membranes. Membranes were blocked in Tris-buffered saline (TBS) containing 5% non-fat dry milk and 0.1% Tween-20 for 1 hour at room temperature. Membranes were then incubated overnight at 4 °C with primary antibodies diluted in the same blocking buffer at the following concentrations: anti-phospho-TrkA (1:1000; Cell Signaling Technology, Cat. No. 4619), anti-TrkA (1:1000; Cell Signaling Technology, Cat. No. 2510), anti-p75NTR (1:2000; Abcam, Cat. No. Ab52987), anti-RBPMS (1:1000; Abcam, Cat. No. Ab152101), anti-GFAP (1:1000; Abcam, Cat. No. Ab7260), anti-Iba1 (1:5000; Wako Chemicals), and BRN3A (1:500; Santa Cruz Biotechnology, Cat. No. Sc-8429).

Following primary antibody incubation, membranes were washed and incubated with horseradish peroxidase (HRP)-conjugated secondary antibodies (anti-mouse or anti-rabbit; 1:5000; Santa Cruz Biotechnology, Milan, Italy) for 1 hour at room temperature. Immunoreactive bands were visualized using enhanced chemiluminescence (ECL) detection reagents (Amersham, Westar; Cat. No. XLS142, Bologna, Italy). Β-Actin was used as an internal loading control. Band intensities were quantified by densitometry using ImageJ software, and data were derived from a minimum of three independent experiments.

### Immunofluorescence on whole-mount retinae

Whole-mount immunofluorescence was performed to assess retinal cellular populations and TrkA receptor distribution. Following euthanasia, eyes were enucleated and fixed in 4% paraformaldehyde (PFA) prepared in 1× phosphate-buffered saline (PBS; sc-281692, Santa Cruz Biotechnology, Heidelberg, Germany) for 1 hour at room temperature (RT). Post-fixation, eyes were washed in PBS and cryoprotected overnight at 4 °C in 30% sucrose (S0389, Sigma-Aldrich, Darmstadt, Germany) in PBS. Retinas were then dissected as flat mounts with four radial cuts to delineate the retinal quadrants.

Following three 10-minute PBS washes, tissues were incubated overnight at 4 °C in blocking solution comprising 5% normal goat serum (NGS; 566460, Sigma-Aldrich) and 0.5% Triton X-100 (93443, Sigma-Aldrich) in PBS. Samples were then incubated for 5 days at 4 °C in the same blocking solution containing primary antibodies at the following dilutions: RBPMS (1:500; guinea pig; PhosphoSolutions, Aurora, USA; RRID:AB_2492226), MNAC13 (1:500; mouse monoclonal anti-TrkA ^36^, anti-Iba1 (1:1000 guinea pig; #234308 Synaptic System), NeuN (1:1000; chicken; ABN91 Merck), anti-Iba1 (1:500; rabbit; Wako, Richmond, USA; RRID:AB_839504), anti-CD68 (1:300; mouse; Serotec, Neuried, Germany; RRID:AB_2291300), anti-GFAP (1:500; mouse; Sigma-Aldrich), anti-p75NTR #9651(1:1000; rabbit polyclonal antiserum to the extracellular domain of mouse p75 ^37^ (gift of Moses Chao).

On day six, retinae were washed three times for 10 minutes in PBS containing 0.3% Triton X-100 and incubated for 2 days at 4 °C with species-specific secondary antibodies (1:1000 in PBS with 0.3% Triton X-100 and 5% NGS). The following fluorophore-conjugated antibodies were used: Alexa Fluor 647 goat anti-guinea pig (A21450, Thermo Fisher, USA; RRID: AB_2535867), Alexa Fluor 555 goat anti-mouse (A32727, Thermo Fisher; RRID:AB_2633276), and Alexa Fluor 488 goat anti-rabbit (A32731, Thermo Fisher; RRID: AB_2633280), Alexa Fluor 647 anti-gp (A21450 Invitrogen), Alexa Fluor 555 anti-rabbit (A31572 Invitrogen), Alexa Fluor 488 anti-chicken (A11039 Invitrogen).

Subsequently, samples were washed three times in PBS, then counterstained overnight at 4 °C with DAPI (1:1000; D9542, Sigma-Aldrich) in PBS containing 0.3% Triton X-100. After final PBS washes, retinae were mounted onto glass slides using Vectashield mounting medium (H1000, Vector Laboratories, Newark, USA) and coverslipped for imaging.

### Confocal microscopy

Structured illumination microscopy (SIM) was employed for high-resolution imaging of retinal whole mounts using an X-light V2 spinning disk confocal system (CrestOptics, Italy) equipped with a Video Confocal super-resolution module, a 40×/1.25 NA PlanApo Lambda oil immersion objective (Nikon, Japan), a Zyla sCMOS camera (Andor), and a Spectra X Lumencor LED light source with bandpass filters at 460–490 nm and 535–600 nm (Chroma Technology, USA). All fluorescence channels (Alexa Fluor 488, 555, 647, and DAPI) were acquired simultaneously. Raw 16-bit SIM datasets were processed and channel-wise reconstructed using Metamorph software.

### Cell counts

Cell counts were obtained from confocal z-stacks spanning the full depth of the retinal ganglion cell layer (GCL; 10–35 planes per stack). Quantification was performed at predefined retinal locations using 187.5 μm × 187.5 μm fields of view (40× objective with 2× digital zoom). Cells positive for RBPMS and Iba1 were counted within the ganglion cell layer (GCL)-inner plexiform layer (IPL) and outer plexiform layer (OPL). Analysis was conducted systematically from the upper-left to the lower-right of each field.

DAPI staining was used to distinguish intact nuclei from apoptotic profiles. Total cell counts per retina were derived by calculating the mean cell density (cells/mm²) and multiplying by the total retinal area, which was measured using TIFF images analyzed in ImageJ (NIH). All quantifications were performed blind to the experimental groups.

### Microglial Morphological Analysis

Microglial morphology was assessed based on the approach described by Marrone ^38^. Confocal z-stacks encompassing the full retinal thickness were acquired at fixed positions (187.5 μm × 187.5 μm fields; 40× objective; 2× digital zoom). Maximum intensity projections were generated and analyzed using the “Shape Descriptor 1u” plugin in ImageJ (NIH). Images were converted to 8-bit grayscale, and thresholds were manually optimized to isolate cell structures while minimizing artifacts. Morphometric parameters measured included cell area (A, in μm²), perimeter (P, in μm), and cell density. The transformation index (TI), a measure of cell shape complexity, was calculated using the formula: TI = (P²) / (4πA). Lower TI values correspond to amoeboid, less ramified microglia, while higher values indicate increased ramification. To control for cell size, the area-to-transformation index ratio (A/TI) was also computed, with lower A/TI values representing more complex morphologies. All morphological analyses were conducted in a blinded fashion.

### Statistical Analyses

Statistical analyses were conducted using GraphPad Prism 8 (GraphPad Software, San Diego, CA, USA). Data are presented as mean ± standard error of the mean (SEM). For comparisons between two groups, two-tailed unpaired Student’s t-tests were applied. The Shapiro–Wilk test was used to assess normality; non-parametric tests were employed if distribution assumptions were not met. Statistical significance was defined as follows: *P value < 0.05, **P value < 0.01, and ***P value < 0.001.

## Results

### PLX5622 dietary administration effectively depletes retinal microglia in C57BL/6J mice

To evaluate the efficacy of PLX5622 in depleting retinal microglia, we quantified microglial populations in both male and female 2-month-old C57BL/6J mice following dietary administration of PLX5622. Age-and sex-matched mice maintained on standard chow served as controls. After three weeks of PLX5622 treatment, whole-mount retinal immunofluorescence revealed a marked reduction in Iba1⁺ immunoreactivity, a specific marker for microglia (Fig. 1A–B). Quantitative analysis confirmed a significant 77% reduction in Iba1⁺ cell density in PLX5622-treated retinas compared to controls (P < 0.001; Fig. 1C). Layer-specific analysis of microglial distribution further revealed a threefold decrease in Iba1⁺ cell number and microglial coverage area within the GCL-IPL of PLX5622-treated mice relative to controls (P < 0.001; Fig. 1D). In the outer plexiform layer (OPL), PLX5622 treatment also resulted in a substantial decrease in microglial cell number, accompanied by 88% reduction in microglial coverage area (P < 0.001; Fig. 1E). Western blot analysis demonstrated a three-fold reduction in Iba1 protein levels in treated retinas compared to controls (P < 0.001; Fig. 1F).

**Figure 1.**
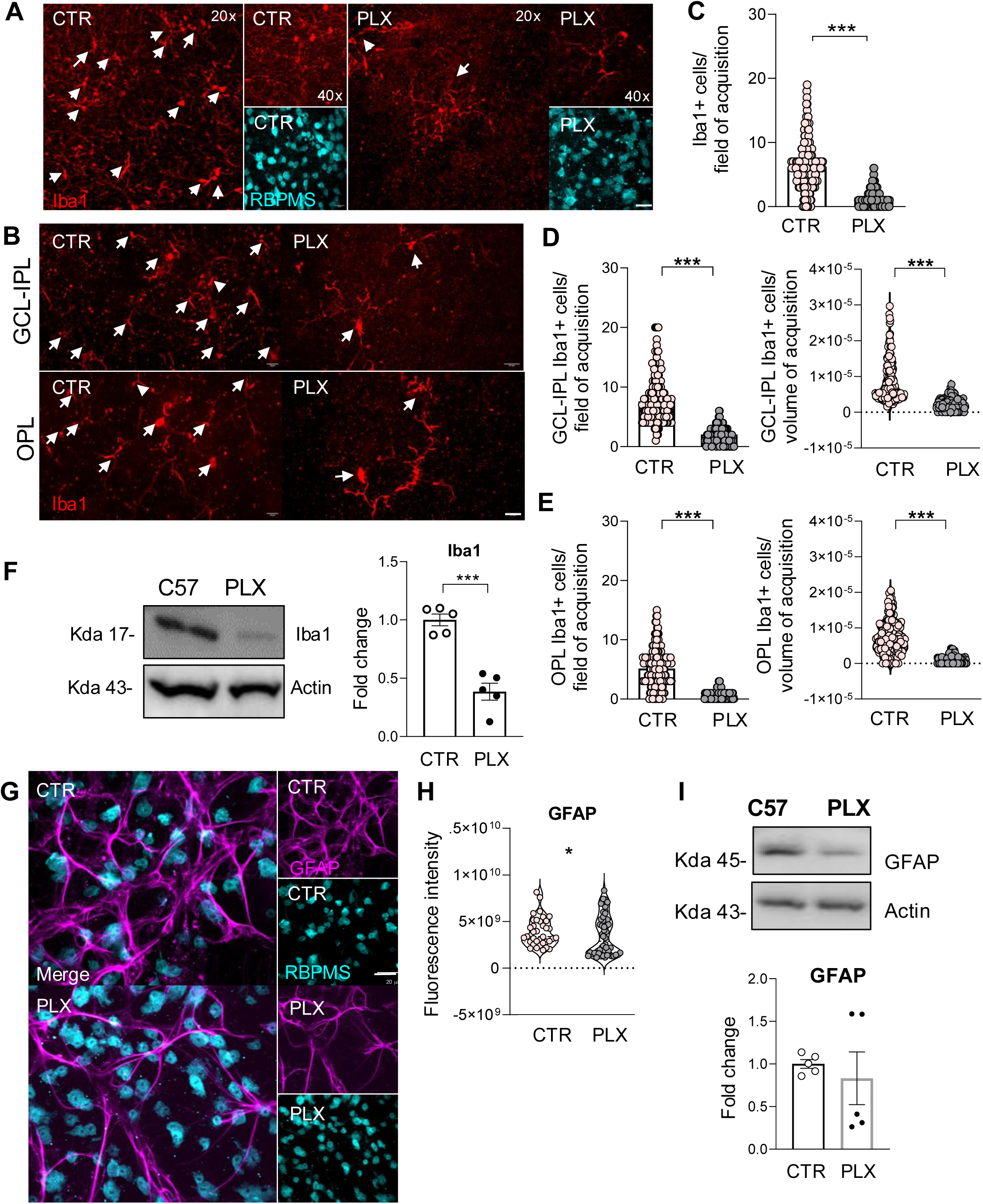
Dietary administration of PLX5622 efficiently depletes retinal microglia in both GCL/IPL and OPL. **A-B)** Representative immunofluorescence images of retinal whole-mounts stained for Iba1 (microglia, red) and RBPMS (retinal ganglion cells, cyan), showing substantial microglial depletion in mice fed PLX5622-formulated chow compared to controls. **A)** Overview images; scale bar, 20 μm. **B)** High-magnification images (60×); scale bar, 40 μm. **C-E)** Quantification of Iba1+ cell number (**C**) and density (**D-E**) demonstrate significant microglial loss in both the ganglion cell layer (GCL) and outer plexiform layer (OPL) following PLX5622 treatment. **F**) Western blot analysis and relative densitometric quantification of retinal lysates showed a reduction of Iba1 protein level in PLX-treated retinae compared to controls. **G-H**) Representative immunofluorescence images (**G**) and corresponding quantification (**H**) of GFAP+ astrocytes (magenta) and RBPMS+ retinal ganglion cells (cyan) reveal a significant reduction in astrogliosis in PLX5622-treated retinae compared to controls. **I**) Western blot analysis and relative densitometric quantification of retinal lysates showed a reduction of GFAP protein level in PLX-treated retinae compared to controls. Data are showed as Mean ± SEM; n=9 for each experimental group. Statistical significance was assessed using unpaired Student’s t-test: ***P value < 0.001 ctr vs PLX.

Together, these findings demonstrate that dietary administration of PLX5622 for three weeks induces robust and widespread retinal microglia depletion in young adult C57BL/6J mice. The differential reduction across retinal layers suggests potential layer-specific microglial morphological or functional adaptations following depletion.

To test whether microglial depletion affects astrocyte integrity, whole-mount retinas from PLX5622-treated and control mice were immune-stained for the astrocyte marker GFAP (Fig. 1G). Quantitative analysis revealed a significant reduction in GFAP-positive staining in microglia-depleted retinas (P< 0.05; Fig. 1H), suggesting that the absence of microglia also affects the astroglial population. Western blot analysis further supported this observation, revealing a downward trend in GFAP protein levels within PLX-treated retinas (Fig. 1I).

### CSF1R inhibition leads to layer-specific morphological modifications in the residual microglia

Given the well-established relationship between microglial density, morphology, and functional state, we analyzed microglial morphology in retinas from control and PLX5622-treated mice. Whole-mount retinal immunofluorescence revealed that CSF1R inhibition altered the morphology of remaining microglia (Fig. 2A), as further shown by the silhouette used for transformation index (TI) quantification (Fig. 2B)

**Figure 2.**
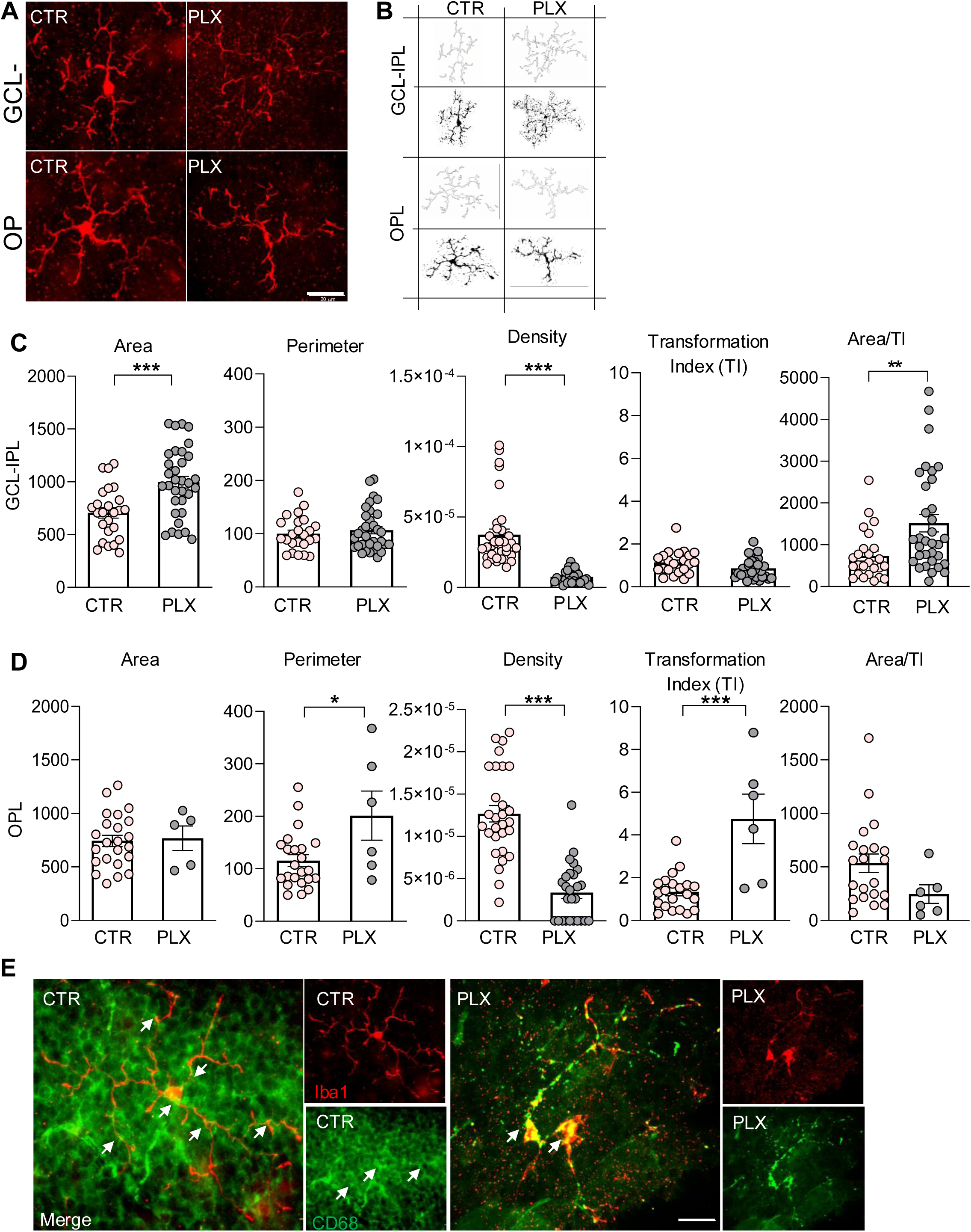
PLX5622 alters the morphology of residual retinal microglia, promoting a more ramified phenotype. **A)** Representative tracings illustrating distinct microglial morphologies observed in control and PLX5622-treated retinae. **B)** Immunofluorescence images showing Iba1+ microglia (red) in the ganglion cell layer-inner plexiform layer (GCL-IPL) and outer plexiform layer (OPL) of control and PLX5622-treated retinae. Scale bar, 20 μm. **C)** Graphical summary of morphological parameters used for analysis (cell area, perimeter, density, transformation index [TI] and ratio between area and transformation index [TI]). Quantitative comparison shows that residual microglia in PLX5622-treated retinae exhibit significantly increased cell area, reduced density and augmented Area/TI in the GCL-IPL, along with increased perimeter and TI in the OPL, indicative of a more ramified morphology. **D)** Representative confocal images showing co-labelling of Iba1 (red) and CD68 (green) in control and PLX5622-treated retinae. Merged images reveal areas of co-localization (yellow), suggestive of altered microglial activation. Scale bar, 40 μm. Data are presented as mean ± SEM (n = 9 per group). Statistical significance was determined by unpaired two-tailed Student’s t-test: *P < 0.05, **P < 0.01, ***P < 0.001 (control vs. PLX5622).

In GCL-IPL, PLX5622-treated retinas exhibited a 41% increase in average microglial cell area and a twofold reduction in microglial density compared to controls (P< 0.001; Fig. 2C). A corresponding decrease in TI values suggested a shift toward a more reactive, likely phagocytic, microglial phenotype. However, we did not observe TI changes in the GCL-IPL microglia. To further assess morphological changes in relation to functional status, we computed the area-to-transformation index (A/TI) ratio. This parameter was significantly reduced in PLX5622-treated retinas, consistent with a transition to a less branched and more compact morphology typical of reactive microglia (P< 0.001; Fig. 2C). In contrast, analysis of the outer plexiform layer (OPL) revealed a 74% increase in microglial perimeter (P < 0.05; Fig. 2D) alongside a 73% reduction in cell density (P < 0.001; Fig. 2D) following PLX5622 treatment. Notably, both TI and A/TI values significantly increased in retinas from PLX-treated mice (P < 0.01; Fig. 2D) indicating a more ramified morphology consistent with a less reactive microglia state in this layer. To further confirm microglial reactive phenotype status, we performed immunofluorescence staining for CD68, a lysosomal marker specifically upregulated in phagocytic microglia. In PLX5622-treated retinas, we observed strong colocalization of CD68 and Iba1 in both soma and processes of residual microglia in GCL-IPL (Fig. 2E), indicative of phagocytic activation.

Taken together, these data demonstrate that CSF1R inhibition via PLX5622 not only reduces microglial density but also induces distinct, layer-specific morphological and functional changes of remaining microglia. While GCL-IPL residual microglia display features consistent with reactive, phagocytic phenotype, in the OPL microglia adopt a more ramified, surveillant morphology.

### Microglial depletion impairs RGC number and retinal integrity

Previous studies have shown that the effects of microglia depletion on RGC vary depending on whether the retina is healthy or injured, and at what developmental stage the depletion occurs ^30,39–42^. To assess how microglial loss influences RGC survival in our model, whole-mount retinas from mice on PLX5622-containing diet or control chow were immune-stained for the RGC marker RBPMS and microglia marker Iba1 (Fig. 3A–B). Quantitative analysis revealed a significant reduction in the number of RBPMS+ cells from PLX5622-treated retinas compared to controls (P< 0.01; Fig. 3C), aligned with a 24% decrease in fluorescence intensity (P< 0.001; Fig. 3D). Although the total area covered by RGCs remained unchanged, their perimeter was significantly reduced in the PLX5622-treated retinas (P< 0.05; Fig. 3F), indicative of a reduced cell size. PLX5622 treatment also led to a significant reduction in RGC density (P< 0.01; Fig. 3G) and a pronounced thinning of the retinal layers (P< 0.001; Fig. 3H). These findings were further supported by western blot, showing a 38% decrease in RBPMs protein levels in PLX5622-treated retinas compared with controls (P< 0.05; Fig. 3I). Analysis of an additional neuronal marker, Brna3, revealed a trend toward decreased expression in PLX5622-treated retinas, though without statistical significance (Fig. 3L). Indeed, Brna3 labels RGCs as well as other neuronal cell types within the retinal layers. Together, these findings indicate that microglial depletion induced by PLX5622 compromises RGC wellness and overall retinal integrity.

**Figure 3.**
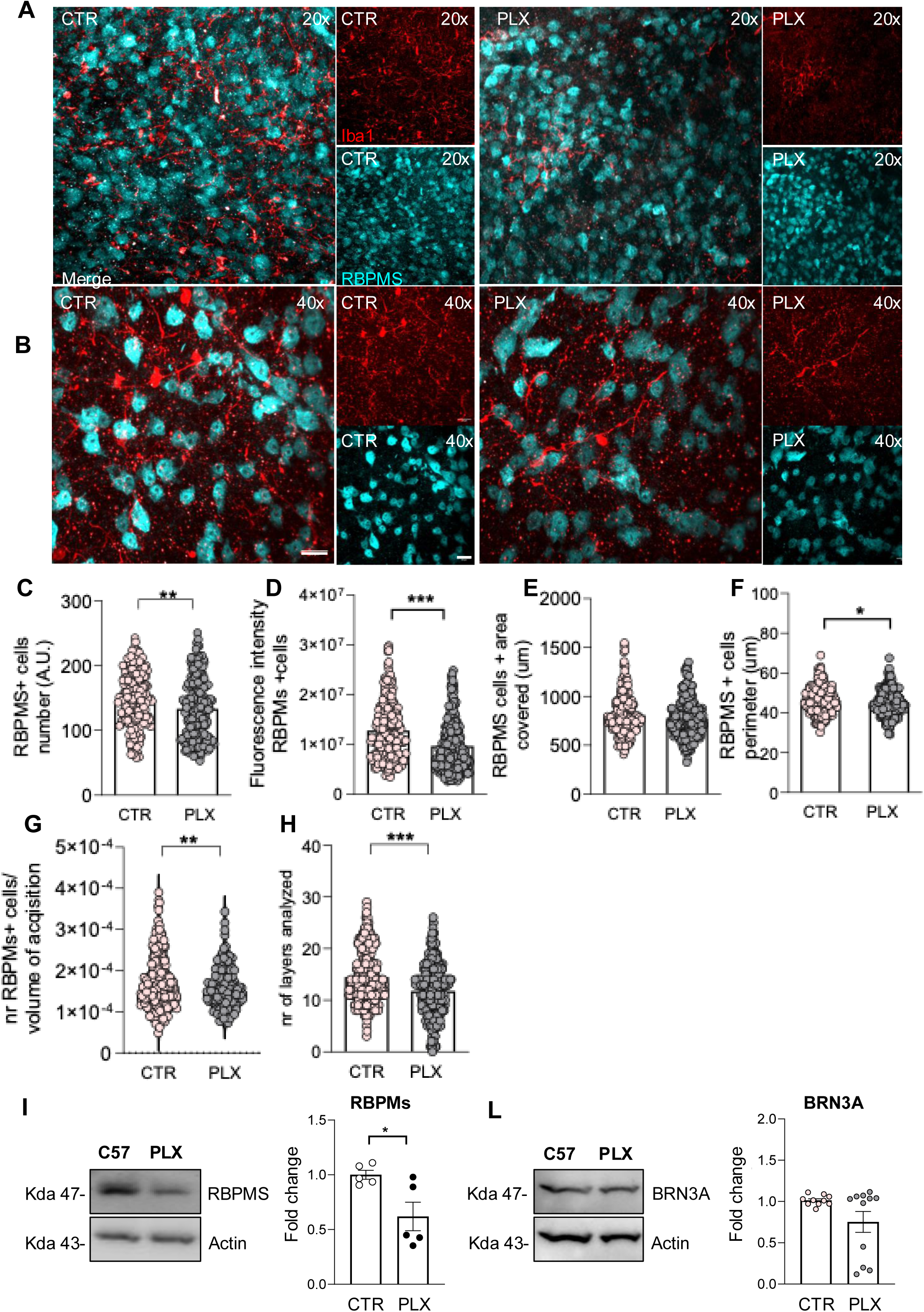
Microglial depletion RGC survival and retinal health. **A–B)** Representative immunofluorescence images of whole-mount retinae stained for RBPMS (a marker of retinal ganglion cells, labelled in cyan) and Iba1 (a microglia marker, labelled in red) from mice fed a PLX5622-containing diet compared to controls. **A)** Overview images; scale bar: 10 μm. **B)** High-magnification views acquired at 40×; scale bar: 20 μm. **C-H)** Quantitative analysis of RBPMS + RGCs shows a significant reduction in RGC number (**C**), fluorescence intensity (**D**), area covered by RGCs (**E**), RGCs perimeter (**F**), and RGCs density (**G**) in PLX-treated retinae compared to controls. Microglial depletion was also associated with thinning of retinal layers (**H**), indicating alterations in retinal structure. Data are showed as Mean ± SEM; n=9 for each experimental group. Statistical significance was assessed using unpaired Student’s t-test: *P < 0.05, **P value < 0.01, and ***P value < 0.001 ctr vs PLX. **I-L**) Western blot analysis and relative densitometric quantification of retinal lysates showed a reduction of RBPMs and BRN3A protein level in PLX-treated retinae compared to controls. Data are showed as Mean ± SEM; n=5 for each experimental group and each dot represents a pool of three different retinas. Statistical significance was assessed using unpaired Student’s t-test: *P < 0.05 and ***P value < 0.001 ctr vs PLX.

### NGF signalling is downregulated in microglia-depleted retinas

Our previous findings suggested that NGF-induced neuroprotection affects microglia function and morphology by activating TrkA signalling. Indeed, we showed that the TrkA receptor is mainly expressed by microglia. Consequently, NGF-mediated neuroprotective effects are mediated primarily through modulation of microglial function ^34,40,43–46^. To directly verify this mechanism, we examined the expression of NGF receptors TrkA and p75NTR, in retinas from mice receiving PLX5622 and fed with standard diet. Western blot analysis revealed a five-fold reduction in both phosphorylated and total TrkA protein levels in microglia-depleted retinas compared with controls (P< 0.001; Fig. 4D). Notably, TrkA pathway activation by endogenous NGF, assessed by p-TrkA/TrkA ratio, was reduced approximately by 20-fold (P< 0.001; Fig. 4A). Immunofluorescence analysis further confirmed TrkA expression in microglia of control retinas and in the few remaining immune cells of PLX-treated retinas, although at markedly lower (Fig. 4B). In parallel, a significant reduction of p75NTR protein levels was also detected in PLX-treated retinas compared to controls (P < 0.01; Fig. 4A). Microglia reduction was significantly correlated with decreased TrkA expression (P <0.05, r=-0.88, Fig. 4B). In contrast, reduced TrkA signalling in microglia-depleted retinas did not correlate with RGC reduction across the same samples (P = 0.66; r = -0.27; Fig. 4C). Similarly, reduced p75NTR levels did not correlate with the extent of microglia reduction (P=0.85, r=-0.11; Fig. 4D). Immunofluorescence assessment of p75NTR distribution indicated that this receptor was neither expressed by Iba1 positive cells, nor by RGCs (Fig. 4E). Collectively, these findings highlight the essential role of microglia as key and predominant cellular mediators of NGF/TrkA signalling in the retina.

**Figure 4.**
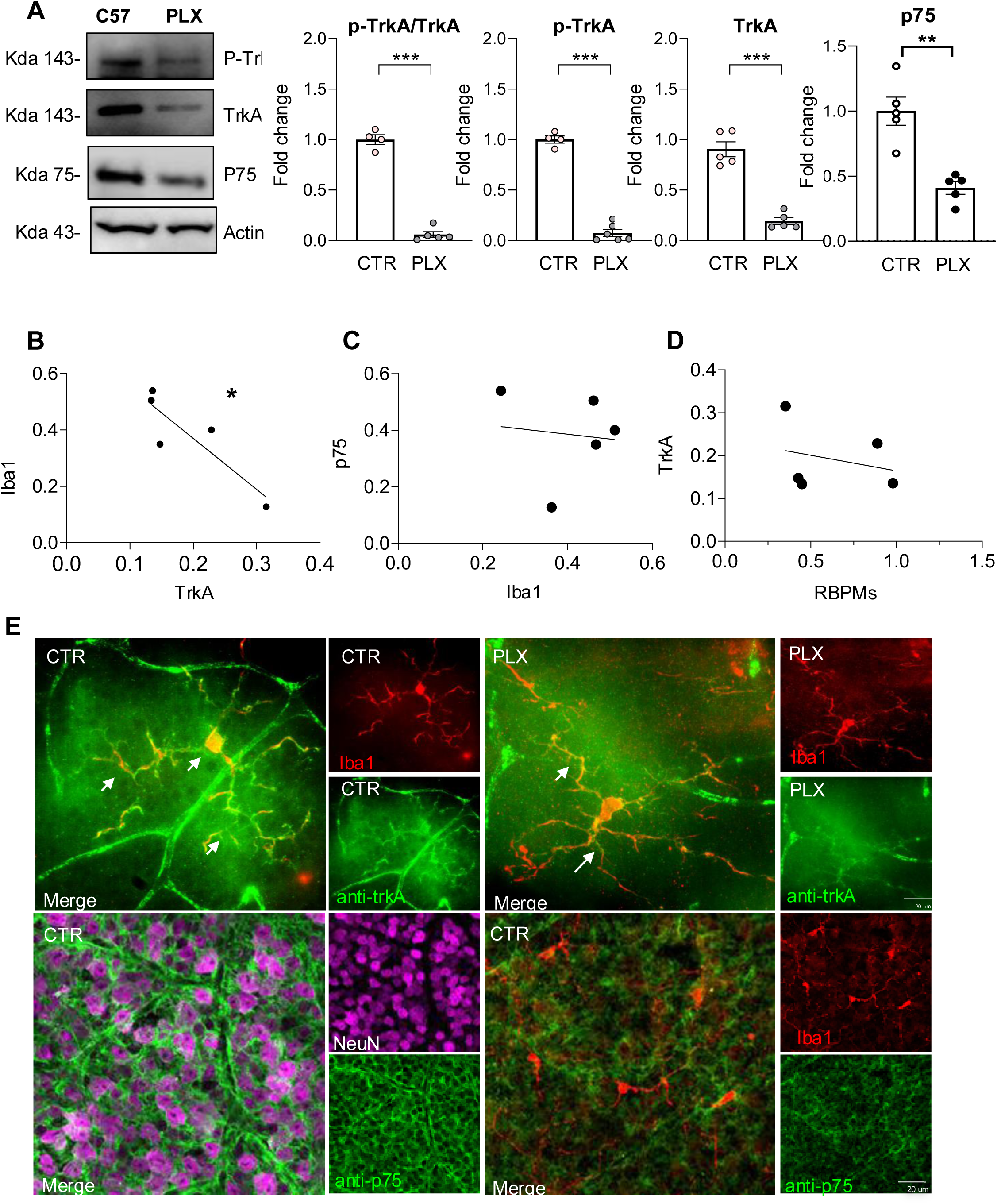
PLX5622 suppresses NGF-TrkA signaling. **A**) Western blot analysis and quantification of retinal lysates showed a strong reduction of TrkA activation, expressed as p-TrkA/TrkA ratio, as well as phosphorylated TrkA and total TrkA and p75NTR protein levels in PLX treated mice compared to the control. **B-D**) Microglia and TrkA activation was strictly correlated in retinae derived from mice fed a PLX5622. Scatter plot showing the relationship between the Iba1 and TrkA (B) protein expression level in retinae derived from mice fed a PLX5622 (r = 0.8788, P < 0.05). In contrast, Iba1 and p75 as well as TrkA and RBPMs protein expression level were not correlated. Pearson correlation’s coefficient was used for correlation analysis. **E**) Confocal images of retinal sections co-immunolabelled for TrkA (green) and Iba1 (red) showed TrkA–Iba1 co-localization (yellow) in control retinae, which is diminished following PLX5622 treatment, indicating that TrkA signaling is largely restricted to microglia. **F**) Confocal images of retinal sections co-immunolabelled for anti-p75 (green), NeuN (magenta) and Iba1 (red) showed p75–Iba1 co-localization (yellow) and perinuclear localization of p75 on RGCs in control retinae, indicating that p75 signaling is located on both microglia and RGCs. Scale bar, 20 μm. Data are presented as mean ± SEM; n = 5 per group, with each data point representing pooled lysates from 3 retinae of different mice. Statistical analysis was performed using unpaired two-tailed Student’s t-test: *P < 0.05, **P < 0.01, ***P < 0.001 (control vs. PLX5622).

## Discussion

In the present study, we provide evidence that NGF-TrkA signaling in the adult mouse retina critically depends on the presence of microglia. Using pharmacological depletion of microglia via CSF1R inhibition, we demonstrate a marked reduction in both total and phosphorylated TrkA levels, accompanied by an impaired RGC survival and alterations in retinal structure. These findings support microglia as key cellular mediators of NGF-dependent neuroprotective signaling in the retina. NGF signaling is classically associated with neuronal survival and differentiation through activation of TrkA and its downstream pathways, including PI3K/Akt and MAPK/ERK cascades ^47–49^. In parallel, the low-affinity receptor p75NTR modulates NGF responses in a context-dependent manner, contributing either to survival or apoptotic signaling depending on receptor co-expression and ligand availability ^50,51^. In the retina, NGF has been shown to promote RGC survival and to exert protective effects in models of neurodegeneration, including glaucoma and optic nerve injury ^34,52–57^. However, it is still not entirely clear which specific cells in the retina are the main targets for NGF. Our findings indicate that microglia represent a major cellular target of NGF signaling, functioning as the bridge that turns neurotrophic instructions into neuroprotective outcomes. Consistent with our previous studies, we confirm that TrkA is predominantly expressed in retinal microglia, with more limited expression in neuronal populations ^34^. Importantly, microglial depletion resulted in a near-complete loss of TrkA protein, strongly suggesting that the majority of NGF-TrkA signaling in the retina is mediated through microglial cells. This interpretation is further supported by the significant correlation between microglial density and TrkA expression. In contrast, the lack of correlation between TrkA levels and RGC number suggests that NGF signaling may not act directly on RGCs but instead exerts its neuroprotective effects indirectly through modulation of microglial function.

This is in line with the emerging view that microglia regulate neuronal survival by integrating environmental signals and releasing trophic or inflammatory mediators ^58,59^. The observed reduction in p75NTR expression following microglial depletion further suggests that NGF signaling is globally altered in the retina under these conditions. Although in our analysis p75NTR was not detected in microglia or RGCs and its reduction does not correlate with microglia levels, its downregulation may reflect changes in other retinal cell types or a general disruption of neurotrophin signaling balance. Given the complex interplay between TrkA and p75NTR, further studies will be required to clarify the contribution of this receptor to retinal homeostasis and degeneration ^51,60^.

The functional consequences of impaired NGF signaling following microglial depletion are reflected in the reduction of RGC number and the overall thinning of retinal layers observed in our study. Loss of NGF-TrkA signaling may reduce trophic support, alter microglial homeostatic functions, and compromise their ability to maintain neuronal integrity. In addition, disruption of NGF signaling may shift microglia toward dysfunctional or maladaptive states, further contributing to neuronal vulnerability. Notably, RGC loss was observed in the absence of any lesion or insult, besides microglia depletion. Although RGC loss did not directly correlate with TrkA levels in our dataset, this may reflect the involvement of additional signaling pathways or compensatory mechanisms, as well as potential threshold effects in neurodegeneration. Together, these findings support a model in which NGF signaling contributes to retinal homeostasis primarily through microglia-dependent mechanisms rather than direct neuron-autonomous effects. We also demonstrate the pivotal role of microglia in supporting RGC survival and maintaining the structural integrity of all layers in young-adult retinas of C57BL/6J mice. The impact of microglial depletion on RGC survival in C57BL/6 control mice varies across developmental stages.

During embryonic development, microglial depletion increases RGC density without altering apoptosis, indicating that microglia eliminate viable RGCs via phagocytosis rather than by inducing cell death ^39^. In contrast, no effect is observed in young adult mice when RGCs are assessed by Brn3a immunolabeling ^42,54^. However, Brn3a lacks specificity, as it also labels other neurons in the ganglion cell layer (GCL), such as amacrine cells. Thus, unchanged Brn3a-positive cell numbers do not exclude RGC alterations. Consistently, in our study, Brn3a immunoreactivity was unchanged following microglial depletion, whereas RBPMS-positive RGCs differed between conditions, suggesting a supportive role for microglia in maintaining neuronal viability under homeostatic conditions. Within this context, NGF-TrkA signaling may represent a key mechanism through which microglia exert neuroprotective effects. In addition to changes in cell number, microglial depletion induced marked alterations in the morphology and functional state of the remaining microglial population. Residual microglia exhibited layer-specific phenotypes, characterized by a more amoeboid, CD68-positive, phagocytic profile in the GCL/IPL and a more ramified morphology in the OPL. These findings indicate that CSF1R inhibition not only reduces microglial density but also reshapes the functional landscape of the remaining cells, potentially reflecting compensatory or repopulation-associated processes. The increased colocalization of Iba1 and CD68 further supports the notion that surviving microglia adopt a reactive, phagocytic phenotype, consistent with previous reports describing functional reprogramming of residual microglia following depletion ^61^. Importantly, microglial depletion also affected other glial populations, as evidenced by reduced GFAP expression and alterations in astrocyte integrity.

This observation highlights the existence of a tightly regulated crosstalk between microglia and astrocytes in maintaining retinal homeostasis. Previous studies have shown that microglia influence astrocyte phenotype and function, including the induction of neuroprotective astrocytic states ^62^. Overall, our findings support a model in which microglia serve as central regulators of NGF-dependent signaling in the retina. Within this context, NGF does not act solely through direct neuronal receptors but instead engages microglia to modulate the retinal environment and promote neuronal survival. Consequently, depletion of microglia disrupts this neurotrophic axis, leading to impaired signaling, altered glial interactions, and increased neuronal vulnerability. These results strengthened the concept that microglia exert both protective and detrimental effects depending on context, and that their neuroprotective role is closely linked to their ability to respond to trophic signals such as NGF and TrkA-biased agonist such as the p75NTR-sparing NGF ^34^.

## Acknowledgements

This study was support by private fundings of EBRI Institute Rita Levi Montalcini Foundation. We thank Francesca Malerba and Rita Florio from the EBRI’s Facility NGF Lab for providing the anti-TrkA antibody, MNAC13.

## Abbreviations

(RGCs): Retinal ganglion cells
(GCL): ganglion cell layer
(OPL): outer plexiform layer
(NGF): Neuronal growth factor
(TrkA): Tropomyosin receptor kinase A
(p75NTR): p75NTR neurotrophin receptor
(RBPMS): RNA binding protein with multiple splicing
(BRN3A): Brain-specific homeobox/POU domain protein 3A

